# Identification of novel tetracycline resistance gene *tet*(X14) and its co-occurrence with *tet*(X2) in a tigecycline-resistant and colistin-resistant *Empedobacter stercoris*

**DOI:** 10.1101/2020.06.17.155978

**Authors:** Yingying Cheng, Yong Chen, Yang Liu, Yuqi Guo, Yanzi Zhou, Tingting Xiao, Shuntian Zhang, Hao Xu, Yunbo Chen, Tongling Shan, Yonghong Xiao, Kai Zhou

## Abstract

Tigecycline is one of the last-resort antibiotics to treat severe infections. Recently, tigecycline resistance has sporadically emerged with an increasing trend, and Tet(X) family represents a new resistance mechanism of tigecycline. In this study, a novel chromosome-encoded tigecycline resistance gene, *tet*(X14), was identified in a tigecycline-resistant and colistin-resistant *Empedobacter stercoris* strain ES183 recovered from a pig fecal sample in China. Tet(X14) shows 67.14-96.39% sequence identity to the other variants [Tet(X) to Tet(X13)]. Overexpression of Tet(X14) in *Escherichia coli* confers 16-fold increase in tigecycline MIC (from 0.125 to 2 mg/L), which is lower than that of Tet(X3), Tet(X4) and Tet(X6). Structural modelling predicted that Tet(X14) shared a high homology with the other 12 variants with RMSD value from 0.003 to 0.055, and Tet(X14) can interact with tetracyclines by a similar pattern as the other Tet(X)s. *tet*(X14) and two copies of *tet*(X2) were identified on a genome island with abnormal GC content carried by the chromosome of ES183, and no mobile genetic elements were found surrounding, suggesting that *tet*(X14) might be heterologously obtained by ES183 via recombination. Blasting in Genbank revealed that Tet(X14) was exclusively detected on the chromosome of *Riemerella anatipestifer*, mainly encoded on antimicrobial resistance islands. *E. stercoris* and *R. anatipestifer* belong to the family *Flavobacteriaceae*, suggesting that the members of *Flavobacteriaceae* maybe the major reservoir of *tet*(X14). Our study reports a novel chromosome-encoded tigecycline resistance gene *tet*(X14). The expanded members of Tet(X) family warrants the potential large-scale dissemination and the necessity of continuous surveillance for *tet*(X)-mediated tigecycline resistance.

## Introduction

Antimicrobial resistance (AMR) represents a major global public health challenge in the 21st century [1]. The clinical infections caused by AMR bacteria, especially carbapenem-resistant *Enterobacteriaceae* (CRE) and *Acinetobacter* spp. (CRA), largely limit the effective prevention and treatment strategies resulting in a high mortality [2,3]. Tigecycline, the minocycline derivative 9-tert-butyl-glycylamido-minocycline, is the third generation of tetracycline family antibiotic which negates most tetracyclines resistance mechanisms due to ribosomal protection and drug efflux [4,5]. This expanded spectrum antibiotics approved by US FDA in 2005 can be used to treat multidrug-resistant gram-positive and gram-negative pathogens [6]. Currently, tigecycline is one of last-resort antibiotics frequently used as a major treatment regimen for the infections caused by CRE and CRA.

Tigecycline resistance has emerged in the clinical setting since then and the resistance is frequently caused by the overexpression of non-specific active efflux pumps or mutations within the drug-binding site in the ribosome [7,8]. Additionally, tigecycline resistance can be mediated by a flavin-dependent monooxygenase gene *tet*(X) and its variants in a small proportion of tigecycline-resistant *Enterobacteriaceae* and *Acinetobacter* isolates through the degradation of tigecycline [5]. The tigecycline breakpoint for *Escherichia coli* and *Citrobacter koseri* has been set down from 2 mg/L in version 8 to 0.5 mg/L in version 9 and version 10 by European Committee on Antimicrobial Susceptibility Testing (EUCAST) [9,10].

The *tet*(X) gene was firstly identified in Tn*4351* and Tn*4400* carried by the chromosome of anaerobe *Bacteroides fragilis* [1]. Subsequently, a few chromosome-encoded and plasmid-mediated novel *tet*(X) variants have been found. Chromosome-encoded *tet*(X), *tet*(X1), *tet*(X2), *tet*(X3) and *tet*(X6) have been identified in *Bacteroides fragilis*, *B. thetaiotaomicron*, *Pseudomonas aeruginosa*, *Myroides phaeus*, *Acinetobacter* spp. and *Proteus* spp., mainly isolated from chickens and pigs [11–15]. Plasmid-mediated *tet*(X3), *tet*(X3.2), *tet*(X4), *tet*(X5) and *tet*(X6) have been detected in *A. baumannii*, *Empedobacter brevis*, *E. falsenii*, *A. indicus*, *A. schindleri*, *A. lwoffiiand* and *A. towneri* isolated from chickens, pigs, cattle, shrimp, avian and human [16–22]. Most recently, another 7 variants including *tet*(X7) to *tet*(X13) have been detected in 244 gut-derived metagenomic libraries in America [3]. Of concern, Tet(X4) and Tet(X6) have recently been found to co-exist with *mcr-1* in *E. coli* [24,25]. The convergence of the last-store antibiotic resistance warns the emergence of superbug in the near future.

The rapid emergence of new resistance mechanisms and phenotypes has worsened the current status of AMR controls, and has elevated the public health significance of this issue. Consequently, the identification of novel *tet*(X) variants is important for us to fully understand the landscape of tigecycline resistance mechanism to control its further dissemination. In this study, we reported a novel chromosome-encoded *tet*(X) variant, designated *tet*(X14), in a livestock-associated *E. stercoris* strain.

## Materials and methods

### Bacterial strains

One hundred and twenty nine strains were isolated from stool samples collected from 6 livestock farms in China in 2019. PCR screening of *tet*(X) variants in the collection was performed as previously described [6].

### Antimicrobial susceptibility testing (AST)

AST was performed according to CLSI guidelines (29th edition) [7]. The breakpoints of tigecycline and eravacycline were interpreted according to the recommended points for *E. coli* by EUCAST version 10.0 [10]. *E. coli* strain ATCC25922 was used for the quality control.

### Whole genome sequencing (WGS) and bioinformatic analysis

Total genomic DNA of the tigecycline-resistant isolate was extracted by Puregene Yeast/Bact Kit B (Qiagen, Maryland, US), and was sequenced by using Hiseq 4000 system (Illumina, San Diego, US) and PromethION platform (Nanopore, Oxford, UK). Hybrid assembly was performed by using Unicycler version 0.4.8 [8]. Antibiotic resistance genes were identified by ResFinder 3.2 with identity >90% and coverage >60% [9]. Synteny analysis was performed using Easyfig [10]. Fragments >5 kb that were absent in at least one genome were detected by BLAST and were defined as genomic islands (GEIs) in this study as previously described [1]. Phylogenetic analysis with amino-acid sequences of Tet(X)s was performed by using the maximum likelihood method with default parameters by using Mega X Version 10.0.5 [2]. The amino acid sequences of Tet(X)s were submitted to ESPript 3 server [3] to perform the alignment and predict the secondary structure elements.

### Functional cloning of *tet*(X14)

The fragment from 219 bp upstream to 53 bp downstream of *tet*(X14) including the predicted promoter of *tet*(X14) was amplified using primers pUC19-*tet*(X14)-F (5’-cgctgcagCAAAAGAGCGGGTTAAGTGG-3’) and p-*tet*(X14)-R (5’-cgtctagaTACTTCACCGGCTCTATTGC-3’). The amplicon was ligated into pUC19, and the recombinant plasmid was transformed into *E. coli* DH5α competent cells by α heat shock. Transformants were selected on LB agar plates containing 100 mg/L ampicillin. In parallel, *tet*(X3), *tet*(X4) and *tet*(X6) were cloned into pUC19 as positive controls.

### Structural modelling of Tet(X14)

The amino acid sequences of Tet(X) variants were submitted to SwissModel [4] to construct 3D structures and 4A6N (PDB entry code) was employed as the template [5]. The overlays of these structures and protein-molecule docking were generated by using AutoDock Vina [6]. Totally hydrogenated Tet(X14), tigecycline and tetracycline were used to perform flexible ligand docking in AutoDock Vina with default parameters. The conformation of ligand which is the most similar with its in 4A6N was chose to construct the recipient-ligand complex to predict the binding sites between Tet(X14) and tigecycline or tetracycline.

### *In silico* screening of *tet*(X14) in GenBank

We screened the sequences of *tet*(X14) in GenBank (https://blast.ncbi.nlm.nih.gov/Blast.cgi, accessed by 10 Jun 2020). Matches with >99.74% identity and >97% coverage were retrieved from GenBank. The retrieved sequences with the reference of each *tet*(X) variant were submitted to phylogenetic analysis to confirm the variant types.

### Nucleotide sequence accession numbers

The complete sequences of the chromosome and plasmids (pES183-1,pES183-2 and pES183-3) of strain ES183 have been submitted to GenBank under the accession numbers CP053698-CP053701.

## Results

### A novel tigecycline resistance gene, *tet*(X14), was identified in *E. stercoris*

A tigecycline-resistant strain ES183 recovered from a pig fecal sample obtained in 2019 was positive for *tet*(X) screening. The strain was identified as *E. stercoris* by using 16S rDNA sequencing [99.86% identity to the 16S rRNA gene of *E. stercoris* strain 994B6 12ER2A (accession no. KP119860)]. Strain ES183 was resistant to amikacin (MIC = 32 mg/L), colistin (MIC = 4 mg/L), and all tetracyclines (MIC = 2-128 mg/L) (Table 1).

**Table 1.** MIC values of antibiotics tested in this study

| Strains | MIC (mg/L) |  |  |  |  |  |  |  |  |  |  |  |  |  |  |  |  |
| --- | --- | --- | --- | --- | --- | --- | --- | --- | --- | --- | --- | --- | --- | --- | --- | --- | --- |
|  | CAZ<sup>a</sup> | CRO | FEP | MEM | CIP | LVX | AMK | CSL | COL | TET | TGC | OTC | CTC | DMC | DOX | MIN | ERV |
| ES183 | 1 | 0.5 | 0.25 | 0.125 | 0.5 | 1 | 32 | 1 | 4 | 16 | 2 | 128 | 8 | 8 | 4 | 0.125 | 1 |
| DH5α-pUC19 | - | - | - | - | - | - | - | - | - | 2 | 0.125 | 2 | 4 | 1 | 2 | 2 | 0.06 |
| DH5α-pUC19-*tet*(X14) | - | - | - | - | - | - | - | - | - | 128 | 2 | 64 | 64 | 32 | 32 | 32 | 4 |
| DH5α-pUC19-*tet*(X6) | - | - | - | - | - | - | - | - | - | 128 | 8 | 128 | 128 | 64 | 64 | 16 | 16 |
| DH5α-pUC19-*tet*(X3) | - | - | - | - | - | - | - | - | - | 128 | 16 | 128 | 128 | 64 | 64 | 16 | 32 |
| DH5α-pUC19-*tet*(X4) | - | - | - | - | - | - | - | - | - | 128 | 16 | 128 | 128 | 128 | 64 | 64 | 16 |

Abbreviation: CAZ, Ceftazidime; CRO, Ceftriaxone; FEP, Cefepime; MEM, Meropenem; CIP, Ciprofloxacin; LVX, Levofloxacin; AMK, Amikacin; CSL, Cefoperazone-Sulbactam; COL, Colistin; TET, Tetracycline; TGC, Tigecycline; OTC, Oxytetracycline; CTC, Chlortetracycline; DMC, Demeclocycline; DOX, Doxycycline; MIN, Minocycline; ERV, Eravacycline.

WGS of ES183 was performed to understand the mechanism of resistance to tetracyclines in ES183. Hybrid assembly of short-read and long-read sequencing data generated a 2.82-Mb chromosome with GC content of 31.89% and 3 plasmids: pES183-1 (10,810 bp; GC content of 24.75%), pES183-2 (2,766 bp; GC content of 33.73%) and pES183-3 (4819 bp; GC content of 25.88%). Three resistance genes were detected in ES183, including two copies of *tet*(X2) and a novel *tet*(X) variant with a size of 1167 bp. The novel *tet*(X) gene encoded a 388-aa protein that displayed 67.14-96.39% identity to reported variants [Tet(X) to Tet(X13)] (Figure 1). Phylogenetic analysis showed that the novel Tet(X) variant formed a clade separated from the reported Tet(X) variants (Figure 1). Taken together, the results suggest that a novel member of *tet*(X) family was identified, designated *tet*(X14). We additionally noted that the amino-acid sequence of Tet(X10) is identical to Tet(X2), that of Tet(X13) is different from Tet(X6) with one amino acid (L368S), and that of Tet(X9) differs from Tet(X7.2) with two amino acids (I156L and G177V).

**Figure 1.**
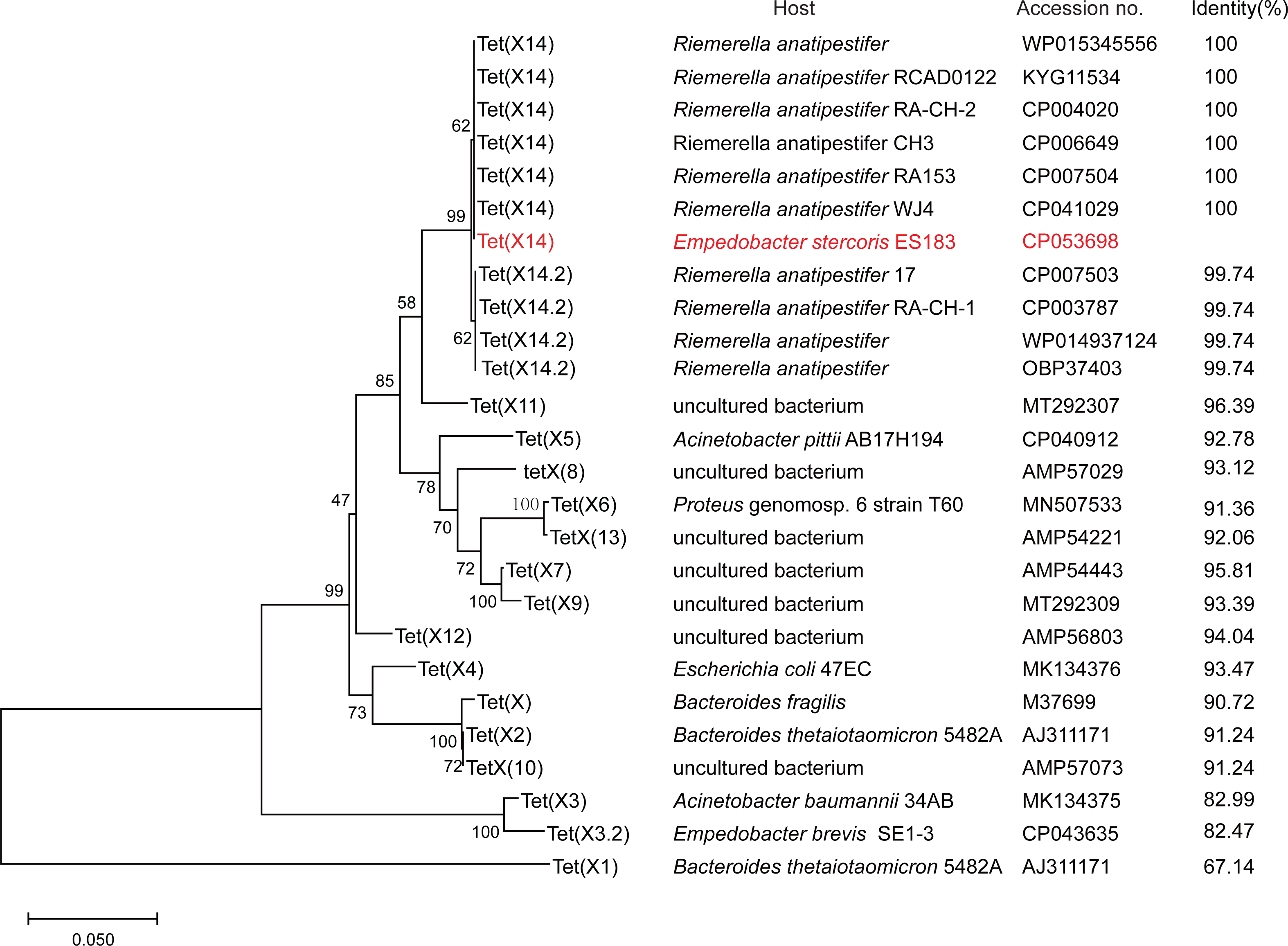
Phylogenetic analysis of the amino acid sequences of Tet(X14) and its homologs. The maximum-likelihood tree was inferred using MEGA X Version 10.0.5 with 1000 bootstraps. Eleven amino acid sequences of Tet(X14) identified in this study and GenBank with the other published Tet(X) variants are included in the analysis. Numbers above each node show the percentage of tree configurations that occurred during 1000 bootstrap trials. The scale bar is in fixed nucleotide substitutions per sequence position. Host strains, accession numbers and identity of each Tet(X) variants relative to Tet(X14) detected in strain ES183 (in red) are listed.

To determine the activity of *tet*(X14) against tetracyclines, the gene was cloned into pUC19 and the resulted recombinant vector was transferred to *E. coli* DH5α to construct the transformant DH5α-pUC19-*tet*(X14). A 16-to 64-fold increase in the MIC of all tested tetracyclines was observed for the *tet*(X14) transformant (Table 1), suggesting that *tet*(X14) was active against tetracyclines. To compare the activity of *tet*(X14) with that of other *tet*(X) variants, we further constructed transformants DH5α-pUC19-*tet*(X3), DH5α-pUC19-*tet*(X4), and DH5α-pUC19-*tet*(X6). The MICs of oxytetracycline, chlortetracycline, demeclocycline, doxycycline and minocycline were comparable among the 4 transformants (2-fold difference), while the MICs of tigecycline and eravacycline were 4-to 8-fold lower for DH5α-pUC19-*tet*(X14) than the other transformants (Table 1). This indicates that Tet(X14) mediated slightly lower level of resistance to tigecycline and eravacycline than Tet(X3), Tet(X4) and Tet(X6).

### Tet(X14) is highly similar with the other Tet(X) variants at the structural level

Alignment of the amino acid sequences of Tet(X14) and the other Tet(X) variants showed that the substrate binding sites and flavin adenine dinucleotide (FAD) binding sites were conserved in all Tet(X) variants with similar secondary structures (Figure S1). The model structure of Tet(X14) was then superposed onto that of other 12 Tet(X) structures [Tet(X2) to Tet(X13)] to perform a homology modelling assay. The overlay of models showed that Tet(X14) shared a high homology with the other Tet(X) variants according to the protein structural architecture (Figure 2A) with RMSD value from 0.003 to 0.055. The data further support that Tet(X14) belongs to Tet(X) family.

**Figure 2.**
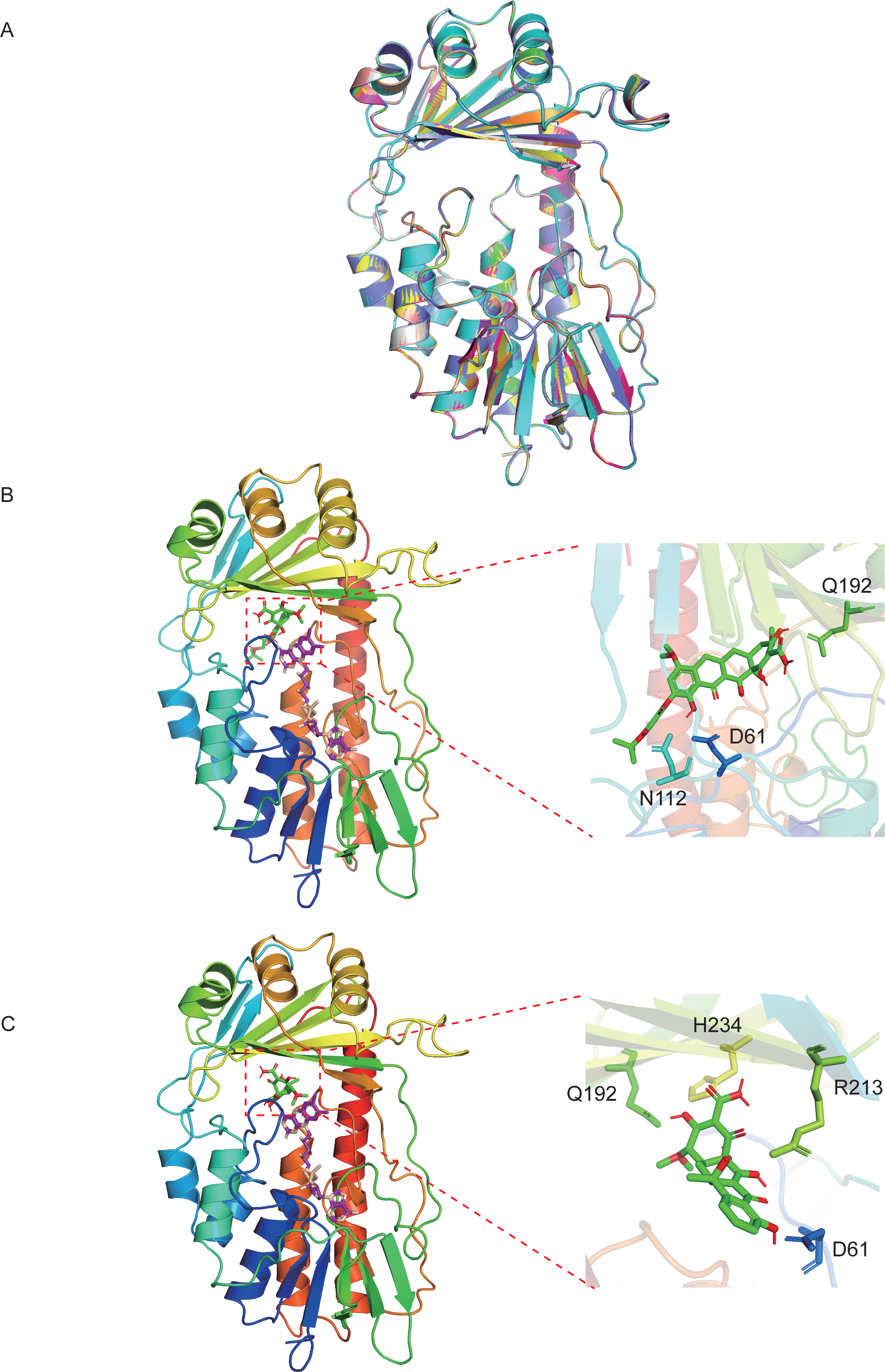
Homology modelling and molecular docking of Tet(X14). (A) Cartoon representation of the modelled Tet(X2) (green), Tet(X3) (cyan), Tet(X4) (magenta), Tet(X5) (yellow), Tet(X6) (pink), Tet(X7) (gray), Tet(X8) (tv_blue), Tet(X9) (orange), Tet(X10) (lime green), Tet(X11) (deep teal), Tet(X12) (hot pink), Tet(X13) (yellow orange) and Tet(X14) (violet purple) structure. Predicted binding conformation of tigecycline (B) and tetracycline (C) (green and red) at the substrate-binding site of the modelled Tet(X14) structure with FAD (violet and wheat). The side chains of residues connected with tigecycline or tetracycline with hydrogen bonds are indicated in the enlarged views.

Flexible ligand docking between Tet(X14) and tetracycline-family antibiotics were performed to predict the hydrogen-bond interaction. D61, N112 and Q192 were the predicted residues involved in interactions between Tet(X14) and tigecycline, and E46, R47, R117 and D311 were the binding sites for FAD cofactor (Figure 2B). This is similar with the structure of TetX2-tigecycline complex derived from the crystallization (PDB no. 4A6N) [5]. Potential interaction sites of Tet(X14) with tetracycline were D61, Q192, H234 and R213 (Figure 2C), which were similar with the modelling of Tet(X6) [5]. These results suggest that as the other Tet(X)s, Tet(X14) interacts with tigecycline and tetracycline through a conserved pattern.

### Tet(X14) was exclusively detected in *Riemerella anatipestifer*

To understand the distribution of Tet(X14) in bacteria, the amino acid sequence of Tet(X14) was blasted in GenBank. Ten hits were obtained with identity >99.74% and coverage >97%, including 4 amino-acid sequences and 6 complete genome sequences (Figure 1). Six hits are identical to the amino acid sequence of *tet*(X14) identified in this study, and the other 4 hits shared 99.74% similarity with only one amino acid difference (G295D), thus designated Tet(X14.2). This is consistent with the phylogenetic analysis that two subclades were formed by Tet(X14) and Tet(X14.2), respectively (Figure 1). The Tet(X14) /Tet(X14.2)-positive isolates exclusively belonged to *R. anatipestifer*. Four Tet(X14) and 2 Tet(X14.2) hits were located on the chromosome of *R. anatipestifer* strains isolated from ducks in eastern and southern China (Table 2). The location of the other 4 hits was undetectable since they were deposited in GenBank as single genes. These results suggest that *R. anatipestifer* might be the major reservoir of Tet(X14).

**Table 2.**
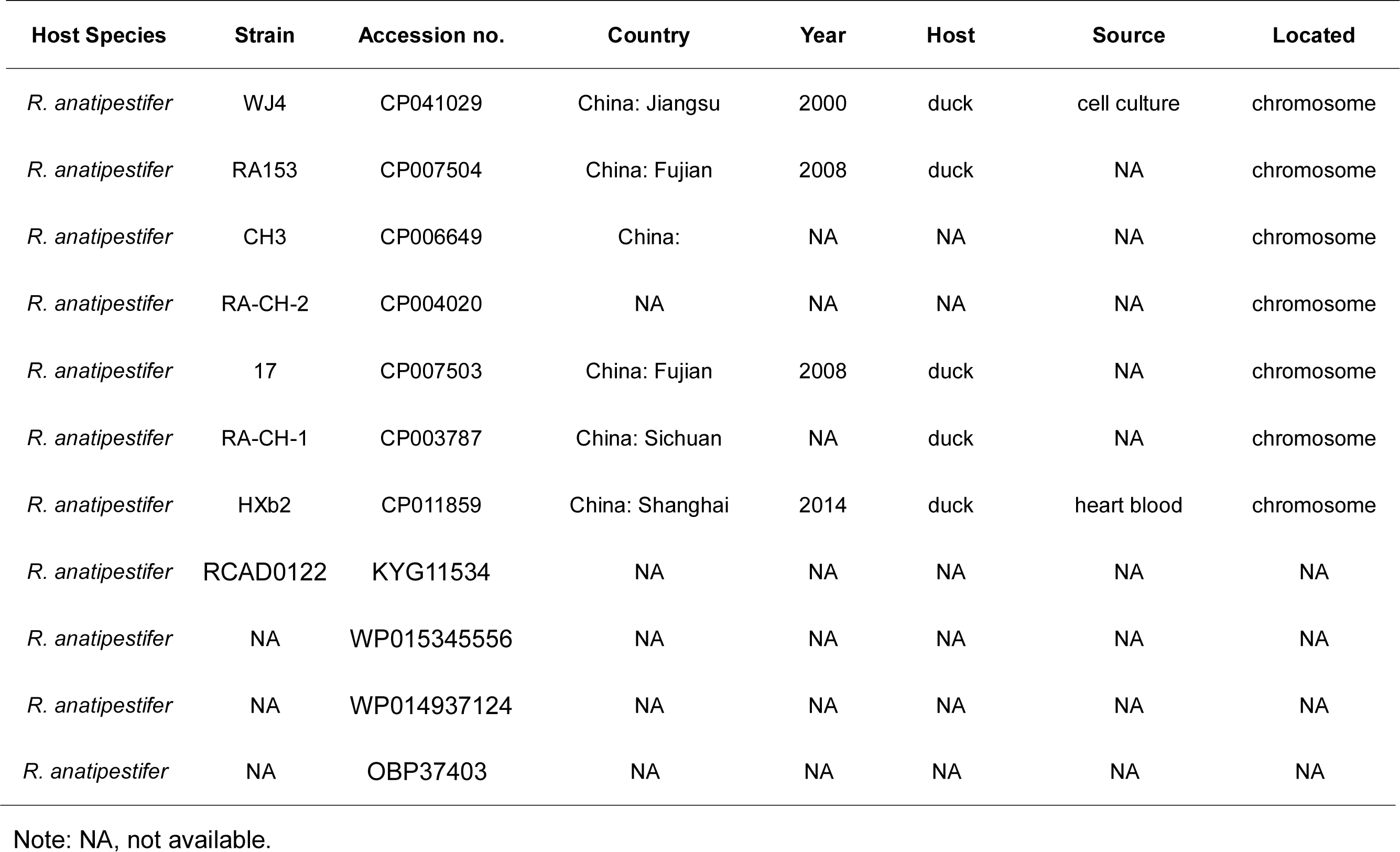
Strains harbouring tet(X14) in GenBank

### Tet(X14) might be obtained by *E. stercoris* strain ES183 via recombination

The *tet*(X14) gene was located at 247668-248834 bp of the chromosome of strain ES183, and an *xerD* gene was found at upstream of *tet*(X14) with opposite direction (Figure 3). It is known that XerD is involved in catalyzing the cutting and rejoining of the bacterial chromosome and plasmid DNA segregation at cell division [37,38]. This adjacency is previously found for plasmid-borne *tet*(X3) and *tet*(X5) [17,19]. No predicted genetic mobile elements, like transposons, integrons or integrative and conjugative elements, were found adjacent to *tet*(X14). To track the source of *tet*(X14) obtained by ES183, the surrounding region of *tet*(X14) (210760-289779-bp) was blasted in GenBank, and two best matches with identity > 86% and coverage > 53% were found, including the chromosome of *E. brevis* BCLYD2 (CP013210) and *E. brevis* SE1-3 (CP043634). The coverage of the other matches was lower than 17%. The fragment (245019-261642-bp) of ES183 encoding *tet*(X14) was not found in *E. brevis* BCLYD2 and *E. brevis* SE1-3, and the flanking regions were conserved in three isolates (Figure 3). The GC content of the *tet*(X14)-encoding region (36.86%) was higher than that of the flanking regions (30.94%-31.71%) and of whole chromosome of ES183 (31.89%). We therefore suppose that the fragment encoding *tet*(X14) is a genomic island (GEI) inserted at the region between genes encoding NUDIX and peptidase M28, which might be obtained from other species via recombination events.

**Figure 3.**
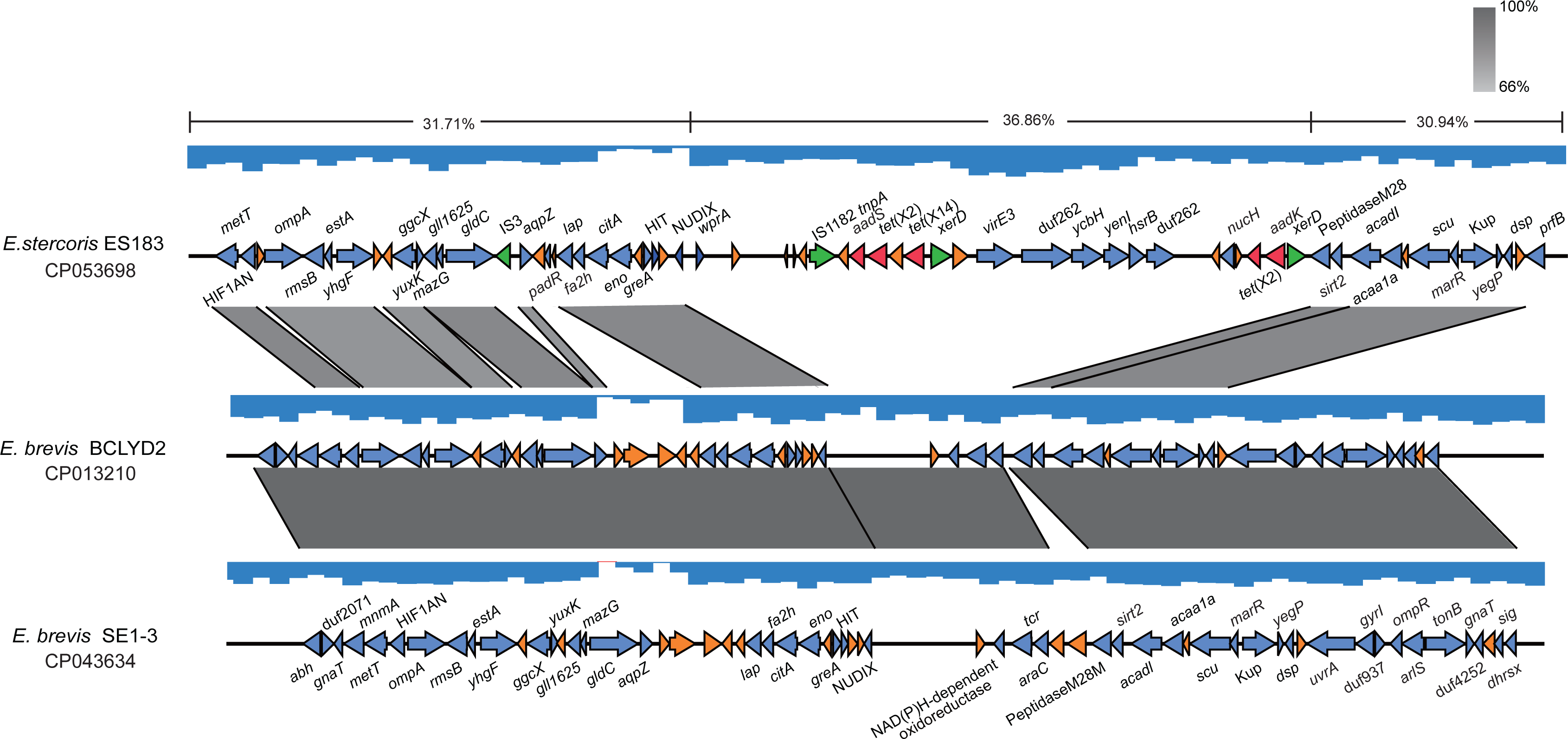
Identification of a genomic island (GEI) encoding *tet*(X14) and *tet*(X2) in ES183 strain. The GEI identified in ES183 inserted between genes encoding NUDIX and peptidase M28. The flanking regions of the GEI are homologous to sequences of two *E. brevis* genomes (CP013210 and CP043634) (>66% identity) retrieved in GenBank shown by grey shading. GC content of the GEI (36.86%) is higher than that of the flanking regions (30.94%- 31.71%) labeled on the top line. The arrows represent the transcriptional direction of the ORFs. Genes are colour-coded, depending on functional annotations: red, antimicrobial resistance; green, mobile genetic elements; blue, other functions; orange, hypothetical proteins.

The surrounding environments of *tet*(X14) identified in *R. anatipestifer* were fully different from that in ES183 (Figure 4). The *xerD* gene was missing in the *tet*(X14) genetic contexts identified in *R. anatipestifer*, and the beta-lactamase gene *bla*OXA-10 adjacent to *tet*(X14) was common. Additionally, two copies of *tet*(X14) with multiple resistance genes were found in most *R. anatipestifer* strains (Figure 4), implying that *tet*(X14) might be encoded on antimicrobial resistance islands (ARIs). Comparative genomics study using *R. anatipestifer* strain ATCC 11845 as the reference identified various *tet*(X14)-encoded ARIs in three genomes of *R. anatipestifer* (CP004020, CP007503, and CP007504) (Figure S2). The *tet*(X14)-encoded ARIs could not be determined in the other genomes due to the lack of suitable reference.

**Figure 4.**
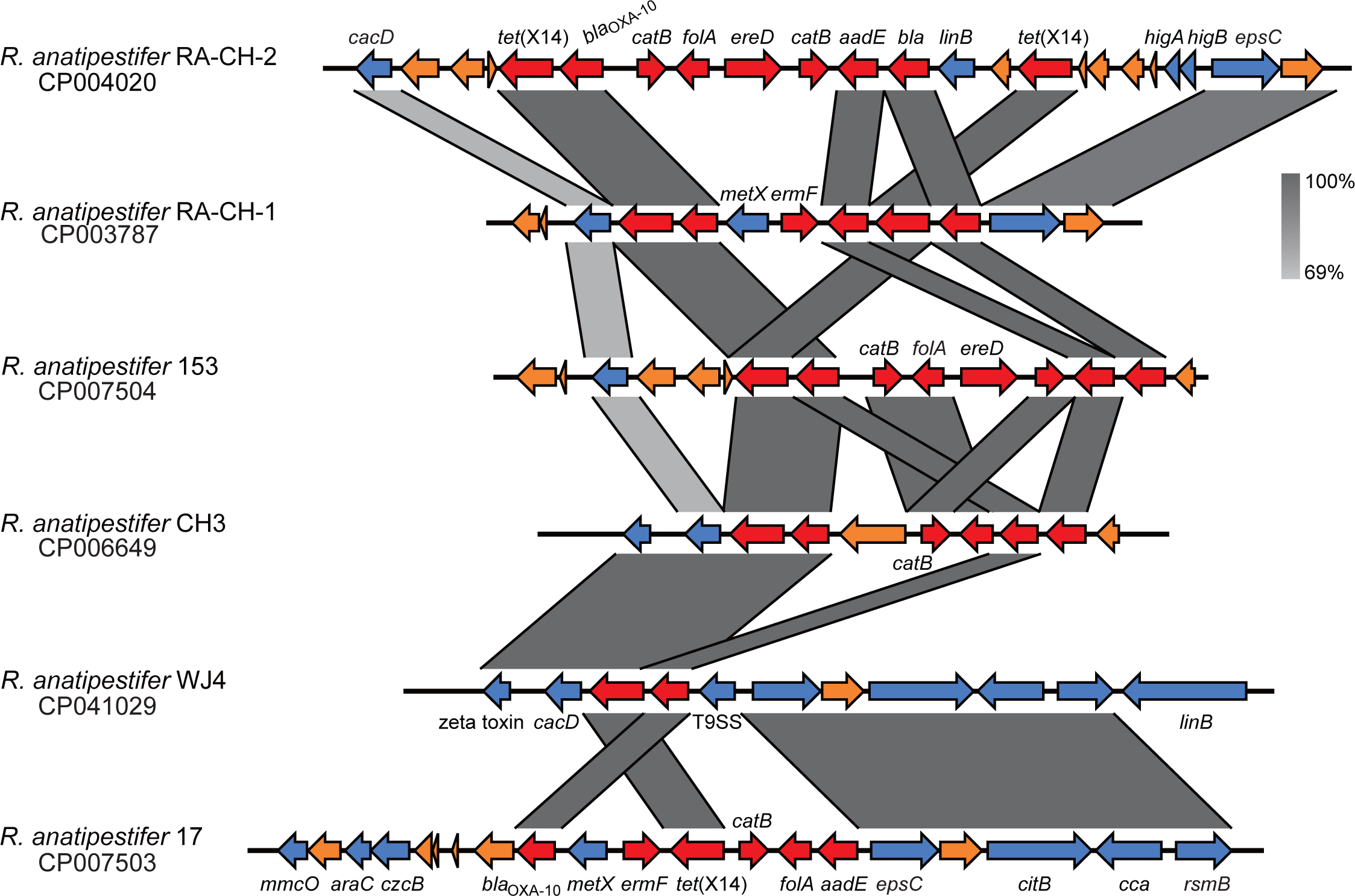
Genomic context of Tet(X14) identified in *R. anatipestifer* strains. The arrows represent the transcriptional direction of the ORFs. Regions of >69% homology are shown by grey shading. Genes are colour-coded, depending on functional annotations: red, antimicrobial resistance; blue, other functions; orange, hypothetical proteins.

Two copies of *tet*(X2) were found at 226213-227379-bp and 249873-251039-bp of the chromosome of strain ES183 with the same transcriptional direction with *tet*(X14). They with *tet*(X14) were located at the same GEI identified above (Figure 3). The two copies of *tet*(X2) were followed by aminoglycosides resistance genes, *aadS* and *aadK*, respectively (Figure 3). This adjacency is similar with that of the firstly reported *tet*(X2) that an *aadS* gene was at upstream of *tet*(X2) in CTnDOT in *B. thetaiotaomicron* 5482A [3]. No Tn structures were found adjacent to *tet*(X2) in strain ES183.

## Discussion

The widespread of CRE represents a large threat to the public health network globally. Currently, tigecycline and colistin are two last-resort antibiotics frequently used to combat lethal infections caused by CRE. However, the wide use of these antibiotics has resulted in the global emergence of resistance in the clinical setting, which significantly compromises their efficacy. Of more concern, the recently identified plasmid-borne resistance genes for colistin (*mcr*) and tigecycline [*tet*(X)] highly challenge the resistance control. At present, the *mcr* variants have already been extended from *mcr*-1 to *mcr*-10, and have disseminated globally [39,40]. It therefore is reasonable to raise the concern that Tet(X) family tigecycline resistance determinants are with potential large-scale dissemination. Indeed, a most recent study dramatically extends the members of Tet(X) family from 7 to 14 [14,15,23]. This study highlights that numerous Tet(X) variants have already circulated in various environmental ecosystem. Monitoring the spread of such resistance genes in the context of One Health (including clinical, animal and environmental sectors) is one of efficient strategies to combat antimicrobial resistance. Identification of new resistance determinants is crucial for fulfilling the strategies and can further aid to improve the current control measures.

In this study, we identified a novel tigecycline resistance gene variant, *tet*(X14), on the chromosome of an *E. stercoris* strain recovered from a pig fecal sample in China. Tet(X14) shows the highest amino acid identity with Tet(X11) (96.39%). Overexpression of *tet*(X14) in *E. coli* DH5α confers 16-fold and 64-fold increase in MIC of tigecycline and eravacycline, respectively. This demonstrated that *tet*(X14) is a novel tigecycline resistance gene. Tet(X14) shows a similar affinity for tetracyclines with the other Tet(X) variants, and their tetracycline-binding sites are conserved. We suppose that the evolutionary pattern of Tet(X) family is of restricted amino acid substitutions with defined limits resulting in a functional consistency against tetracyclines. The resistance activity of Tet(X14) against tigecycline is lower than that of plasmid-mediated Tet(X3), Tet(X4) and Tet(X6) (Table 1). In the crystallization complex of tigecycline and Tet(X2), Q192 and R213 are identified as the common binding sites in four monomers [5]. These two binding sites are conserved in Tet(X4), while hydrogen bonds have been predicted at R211 but not at Q in Tet(X3). This is consistent with the higher tigecycline resistance conferred by Tet(X4) than Tet(X3) in a mouse model [7]. Only Q192 but no any R was identified in the docking complex of tigecycline and Tet(X14) (Figure 2B). This may explain the lower activity of tigecycline resistance conferred by Tet(X14). However, further study should be performed to validate the prediction results. Of note, Tet(X14) was exclusively identified on the chromosome in this study, Tet(X3) and Tet(X4) were frequently detected on plasmids in various species [17,22,41], and Tet(X6) was almost equally distributed on the chromosome and plasmid based on the data available currently [15,42]. A potential correlation was noted for the tigecycline resistance activity and the location of Tet(X)s that the plasmid-borne Tet(X3) and Tet(X4) showed the highest activity (16 mg/L), and the chromosome-encoding Tet(X14) showed the lowest activity (2 mg/L). The activity of Tet(X6) was between them (8 mg/L). However, more data are needed to determine the correlation in the future.

Tet(X14) was exclusively detected in *R. anatipestifer* through blasting in GenBank (Table 2 and Figure 1). *R. anatipestifer* is a Gram-negative bacterium belonging to the family *Flavobacteriaceae*. Intriguingly, *E. stercoris* also belongs to the same family, suggesting that the members of the family *Flavobacteriaceae*, especially *R. anatipestifer*, might be the major reservoir of *tet*(X14). Moreover, all Tet(X14)-producing strains were exclusively detected in China except for one strain with unknown source (Table 2), indicating that Tet(X14) might emerge locally. Of note, *R. anatipestifer* is an important poultry pathogen which primarily causes infection in domestic ducks [3], and *E. stercoris* is livestock associated, which is firstly isolated from a mixed manure sample [4]. We suppose that the emergence of Tet(X14) could be caused by the heavy utilization of tetracycline antibiotics in the animal feed, like tetracycline, oxytetracycline, chlortetracycline, and doxycycline [9]. It is reasonable to predict that such resistance gene would jump into the clinical setting through the food chain and/or zoonosis with high possibilities in the future.

We determined the genetic context of *tet*(X14) to estimate how the gene was captured by the isolates detected here. The surrounding environments of *tet*(X14) identified here were different from those of the other *tet*(X) variants that no any known mobile genetic elements were found. Moreover, the genetic contexts of *tet*(X14) identified in *R. anatipestifer* were completely different from that identified in *E. stercoris*, suggesting that *tet*(X14) was captured by the members of the family *Flavobacteriaceae* individually, and inter-species transmissions might have not occurred yet. An *xerD* gene was found at the upstream of *tet*(X14) identified in *E. stercoris*. It has been reported that XerD is able to mediate the integration of mobile genetic elements (e.g. phages) into the chromosome via homologous recombination [5]. The gene has frequently been found adjacent to other *tet*(X) variants [17,19], It thus would be interesting to validate whether XerD is involved in the mobilization of *tet*(X)s in the future. Additionally, the *tet*(X14) gene was identified on GEIs in *E. stercoris* and on ARIs in *R. anatipestifer* (Fig. S2), and the GC content of the *tet*(X14)-encoding fragment carried by *E. stercoris* was different from the flanking regions. Together, the data imply the heterologous insertions of *tet*(X14) via recombination events. Currently, the limited genomic data largely impedes us to track the source and origin of *tet*(X14).

In summary, we report the discovery of a novel chromosome-encoding tigecycline resistance gene, *tet*(X14), in a tigecycline-resistant and colistin-resistant *E. stercoris* strain. The convergence of resistance to two last-resort antibiotics would largely threaten the global public health system. Tet(X14) has a similar function and structure to other Tet(X) variants, and confers lower tetracycline/glycylcycline MICs than the plasmid-borne Tet(X)s. Recombination may play an important role in the transmission of *tet*(X14). The expanded members of Tet(X) highlights the potential large-scale dissemination and the necessity of continuous surveillance for *tet*(X)-mediated tigcycline resistance.

## Supporting information

Supplementary Figure 1. The secondary structure of Tet(X14) and its homologs.

Supplementary Figure 2. Antibiotic resistance islands (ARIs) encoding tet(X14) identified in R. anatipestifer.

## Acknowledgments

This work was supported by the National Key Research and Development Program of China (2017YFC1200200), Major Infectious Diseases Such as AIDS and Viral Hepatitis Prevention and Control Technology Major Projects (2018ZX10712-001), the National Natural Science Foundation of China (81702045 and 81902029), and Shenzhen Basic Research projects (JCY20190807144409307 and JCY20190730112444588)

---

**Supplementary Figure 1. The secondary structure of Tet(X14) and its homologs.** The alpha helixe and beta sheet of Tet(X14) and the other variants were predicted as the secondary structure elements. The conserved regions among these variants are indicated by red boxes with white letters. The substrates binding sites and FAD binding sites are indicated by asterisks.

**Supplementary Figure 2. Antibiotic resistance islands (ARIs) encoding *tet*(X14) identified in *R. anatipestifer.*** ARIs encoding *tet*(X14) were identified in three *R. anatipestifer* strains (CP007503, CP007504, and CP004020) using *R. anatipestifer* strain ATCC11845 (NC017045) as the reference. The flanking regions of the ARIs are homologous to sequences of ATCC11845 genome with >83% identity shown by grey shading. The arrows represent the transcriptional direction of the ORFs. Genes are colour-coded, depending on functional annotations: red, antimicrobial resistance; green, mobile elements; blue, other functions; orange, hypothetical proteins.

