## Supplementary figures and images for "Identification of novel tetracycline resistance gene *tet*(X14) and its co-occurrence with *tet*(X2) in a tigecycline-resistant and colistin-resistant *Empedobacter stercoris*"

### Supplementary Figure 1. The secondary structure of Tet(X14) and its homologs.

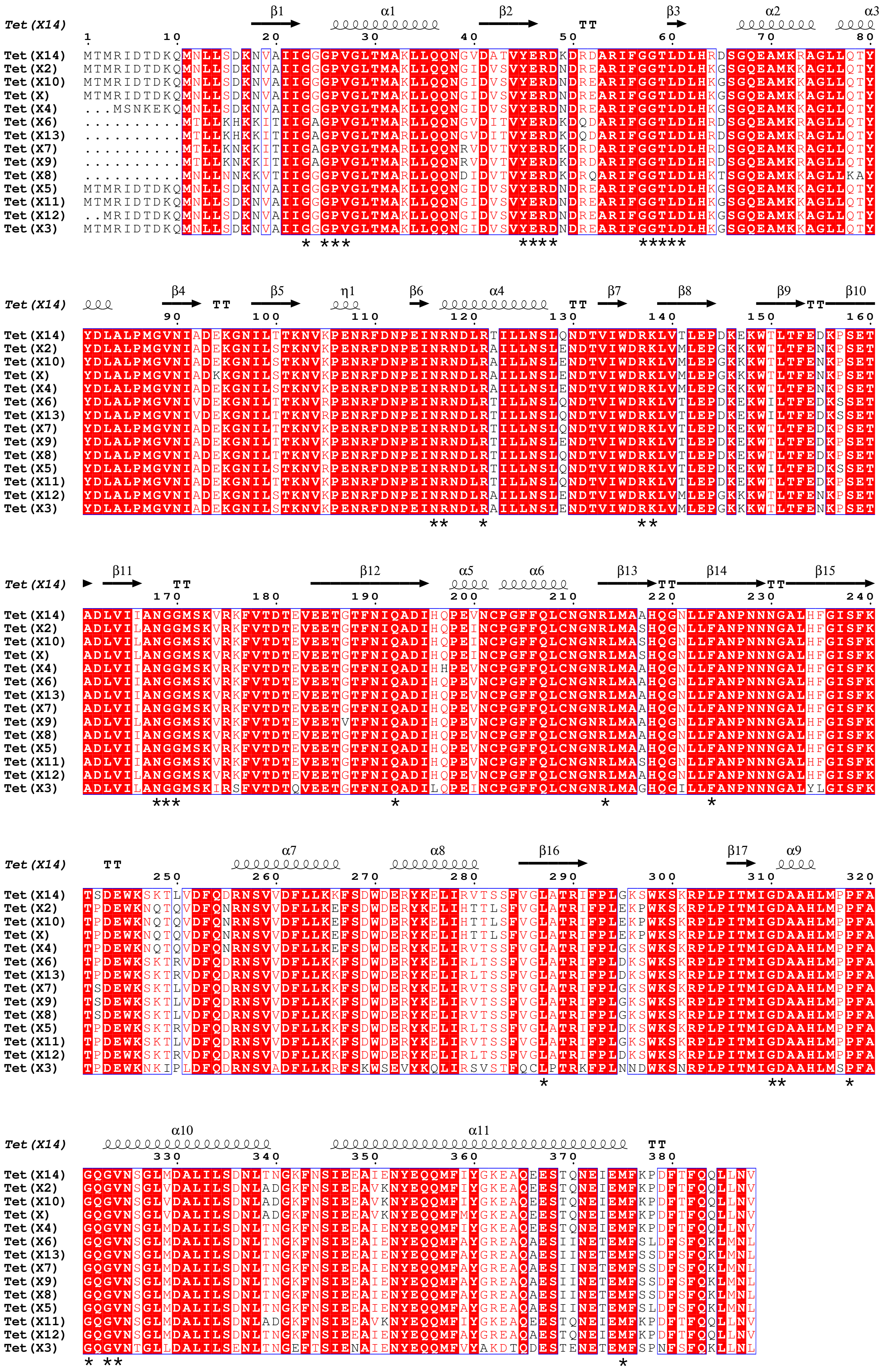

### Supplementary Figure 2. Antibiotic resistance islands (ARIs) encoding tet(X14) identified in R. anatipestifer.

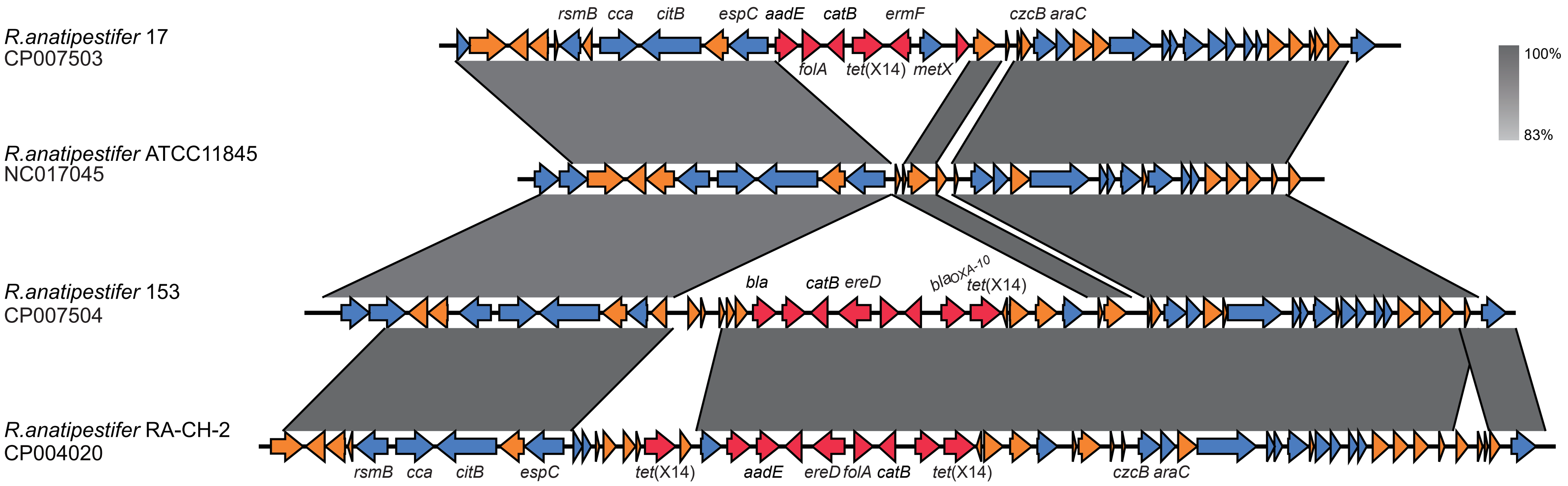
